# Iodine-enhanced micro-CT imaging of soft tissue on the example of peripheral nerve regeneration

**DOI:** 10.1101/477539

**Authors:** Patrick Heimel, Nicole Victoria Swiadek, Paul Slezak, Markus Kerbl, Cornelia Schneider, Sylvia Nürnberger, Heinz Redl, Andreas Herbert Teuschl, David Hercher

## Abstract

Microcomputed tomography (μCT) is widely used for the study of mineralized tissues but a similar use for soft tissues is hindered by their low X-ray attenuation. This limitation can be overcome by the recent development of different staining techniques. Staining with Lugol’s solution, a mixture of one part iodine and two parts potassium iodide in water, stands out among these techniques for its low complexity and cost. Currently, Lugol staining is mostly used for anatomical examination of tissues. In the present study we seek to optimize the quality and reproducibility of the staining for *ex vivo* visualization of soft tissues in the context of a peripheral nerve regeneration model in the rat.

We show that the staining result not only depends on the concentration of the staining solution, but also on the amount of stain in relation to the tissue volume and composition, necessitating careful adaptation of the staining protocol to the respective specimen tissue. This optimization can be simplified by a stepwise staining which we show to yield a similar result compared to staining in a single step. Lugol staining solution results in concentration dependent tissue shrinkage which can be minimized but not eliminated. We compared the shrinkage of tendon, nerve, skeletal muscle, heart, brain and kidney with six iterations of Lugol staining.

60 ml of 0.3% Lugol’s solution per cm^3^ of tissue for 24h yielded good results on the example of a peripheral nerve regeneration model and we were able to show that the regenerating nerve inside a silk fibroin tube can be visualized in 3D using this staining technique. This information helps in deciding the region of interest for histological imaging and provides a 3D context to histological findings. Correlating both imaging modalities has the potential to improve the understanding of the regenerative process.

## Introduction

Microcomputed tomography (μCT) has been used as a valuable tool for understanding and quantifying regenerative processes in bone and other mineralized tissues[1–4]. So far no widespread application of μCT for the evaluation of soft tissues in a similar manner is routinely used due to the low X-ray attenuation of non-mineralized tissues. Commonly used contrast agents such as osmium or gold [5–8] bear the disadvantages of high cost or toxicity. To overcome this limitation, alternative staining methods including iodine-based dyes such as Lugol’s solution were adapted for μCT use, as first described by Metscher[9,10]. Lugol’s solution poses a simple, cost-effective and non-toxic option for contrast enhancement of soft tissues. Yet, its use has been mostly limited to anatomical studies of a wide variety of biological specimen using a broad range of different concentrations of iodine and staining durations, depending on the type of tissue[11]. In regard to soft tissue perfusion, vascular detection in CT scans has been commonly facilitated via contrast agent perfusion. For instance by utilizing intravasal barium sulfate a distinct contrast of blood vessels is achieved[12,13], but incomplete vascular perfusion may lead to an underestimation of the vasculature[14].

The regeneration of peripheral nerve tissue is another field of regenerative medicine which could benefit from a more detailed analysis of e.g. tissue formation or tissue to graft reactions via μCT measurements. Despite growing microsurgical knowledge, the clinical success rate after peripheral nerve repair still remains below 70%[15,16]. Especially defects with a nerve tissue loss larger than 3 cm result in unsatisfying outcomes[17]. Therefore, the treatment of these critical size defects remains an ongoing challenge in peripheral nerve surgeries. In the clinical setting, autologous nerve transplantation is the gold standard for bridging nerve defects. However, the availability of donor nerves is limited and surgery induces an additional loss of function for the patient, resulting in a need for alternative biological or artificial nerve guiding structures to repair these defects[17,18]. So far none of the available guiding structures shows success rates comparable to autologous transplants. Hence there is a strong need for preclinical *in vivo* studies to further characterize and improve such nerve conduits and their effect on regeneration.

Besides functional tests, histological imaging using conventional brightfield microscopy of stained sections is an important tool for evaluation of the regenerating nerve, the respective conduit and interactions between these two structures. A known disadvantage of this method is that slide preparation requires cutting and staining of all samples, which is a destructive and time-consuming process. Therefore, we have established a fast, cost-efficient, reliable and easy-to-perform imaging method based on staining with an aqueous I2KI solution, also known as Lugol’s solution, which enables the generation of 3D images from full size nerve samples to identify specific regions of interest before sectioning. This allows reducing the number of histologic specimens that have to be produced and provides a precise selection of the most suitable region at the same time. Furthermore 3D images allow tracking of the exact route of the supplying vessels, which is important given that vascularization has a strong impact on the regeneration of the peripheral nerve[19,20].

Yu et. al.[21] developed a different technique for simultaneous visualization of blood vessels and nerve tissue. Their method, which employs a barium sulfate perfusion contrast agent to visualize blood vessels, uses Sihler staining[22], which stains the nerves and makes other soft tissue transparent for visualization of nerves. This process is very complex and takes 3 to 4 months from explanting the samples to the completed analysis. Moreover, this imaging technique can only provide 2D images. Three dimensional imaging of nerves in muscles has been shown to be possible with Sihler staining but further increases the complexity of the process and thereby the analysis time[23].

The study presented here seeks to optimize the Lugol staining process for soft tissue regeneration studies by evaluating the effect of different concentrations and a variation in sample storage times before staining. Of special interest to us is to evaluate how samples of unknown tissue composition can be stained in a consistent manner while minimizing the risk of overstaining and other negative side effects, such as shrinkage, of the staining process. For this purpose we stained mouse limbs, which are composed of many different tissues with very similar relative volumes between individuals. Finally the potential of these staining techniques to evaluate peripheral nerve regeneration is demonstrated.

## Material and Methods

### Staining Procedure

Lugol’s solution is one part iodine and two parts potassium iodide in an aqueous solution. To produce a concentration of 0.3%, 0.1% (w/v) iodine and 0.2% (w/v) potassium iodide is dissolved in double-distilled water. All instances of Lugol staining performed in the course of this study were performed in distillated water and mixed according to the ratio above.

### Preparation of Hind Limb, Organ and Tissue Samples

The front and hind limbs of 8 adult mice were used for stain and contrast measurements. Organ and tissue samples of one adult rat for shrinkage measurement included brain, kidney, heart, left and right tibialis, patella and achilles tendons and left and right sciatic nerves. All samples were harvested immediately after euthanasia and fixed in 4% neutral buffered formalin and stored at 4 °C.

After 24 h, the tissue samples were thoroughly rinsed in tap water for at least 1 hour, transferred to 50% ethanol for 1 h and stored in 70% ethanol.

### Preparation of Nerves in a Silk Fibroin Guidance Conduit

Male Sprague–Dawley rats (n=12, Charles River, Germany), weighing between 325 and 415 g, were used in the experiments. Standard rodent food and water were provided ad libitum. All experimental protocols were approved by the City Government of Vienna in accordance with the Austrian law (Animal Use Proposal Permission no. 406382/2015/12) and the National Institute of Health Guidelines.

In this study a silk fibroin based nerve guidance conduit (NGC) described in a previous study[24] was used. Briefly, this NGC is based on a braided tubular structure of raw *Bombyx mori* silk fibers, which has been degummed by boiling in a borate buffer solution[25]. The outside wall is then post-processed via subsequent incubations in the ternary solvent CaCl_2_/H2_O_/ethanol, formic acid and methanol. This combination of treatments results in a fusion of single fibers to a continuous layer which lead to the final favorable mechanical and topographical characteristics of the silk fibroin based NGC.

#### Surgical procedure - peripheral nerve injury

Rats were weighed and anesthetized using isoflurane (induction 3.5%, maintenance 2.0%, Forane^®^) at a flow rate of 800 mL/min. Body temperature was monitored continuously. The right lower limb was shaved and disinfected with povidone-iodine (Betaisodona; Mundipharma) at mid-thigh level. All following surgical procedures were carried out under an surgical microscope (Leica M651; Leica Microsystems). After stump preparation of the muscles, the sciatic nerve was exposed, and a 10 mm segment was excised. A 12 mm silk fibroin conduit as described in[24], was implanted by insertion of the proximal and distal nerve stumps, which were coaptated to the conduit using Ethilon 8/0 epineurial sutures (Ethicon-Johnson and Johnson). Afterwards, the previously dissected muscles were adapted with 4/0 Vicryl^®^ resorbable suture material. Final wound closure was done with the same suture performing intracutane stitches. Analgesic treatment was administered using 0.75 mg/kg bodyweight (BW) meloxicam (Metacam^®^; Boehringer Ingelheim, sid) and 1.25 mg/kg BW butorphanol (Butomidor^®^; Richter Pharma AG, bid) immediately prior to surgical procedure and for 2 days thereafter. After 7 weeks animals underwent perfusion.

#### Barium sulfate perfusion

All rats were perfused with a barium sulfate contrast agent to enable visualization of blood vessels in the defect region as described in [13]. In brief, full body perfusion through the left ventricle with a mixture of heparin and saline was followed by neutral buffered formalin, PBS (phosphate buffered saline, Sigma-Aldrich, St. Louis, USA) and then the contrast agent, pre-heated to 40 °C, in a mixture of 30% barium sulfate (Micropaque, Guerbet, France) and 3% gelatin in PBS. Both the treated and untreated nerves of all rats were harvested. The nerves were kept in 4% neutral buffered formalin at 4°C for 24h. Afterwards the nerves were rinsed under flowing water for at least 1 h to remove formalin, transferred to 50% ethanol for 1 h before being stored in 70% ethanol.

### Staining

#### Hind limb Staining

The mouse hind limbs were stained with different concentrations to determine the effect of iterative staining and to quantify any change of contrast between tissues (Figure 1). They were separated into single and double staining groups, each containing 4 left and 4 right limbs. Each hind limb was stained in 22.5 ml Lugol’s solution in each staining step which was estimated to be approximately 15 ml Lugol’s solution per cm3 of tissue volume. Hind limb Step I: The double staining group was stained twice at 0.15% Lugol’s solution for 72 h each. The single staining group was stained only once with 0.3% Lugol’s solution. Hind limb Steps II/III: The double staining group was stained 4 times with 0.3% Lugol’s solution and the single staining group was stained twice with 0.6% Lugol’s solution.

**Figure 1:**
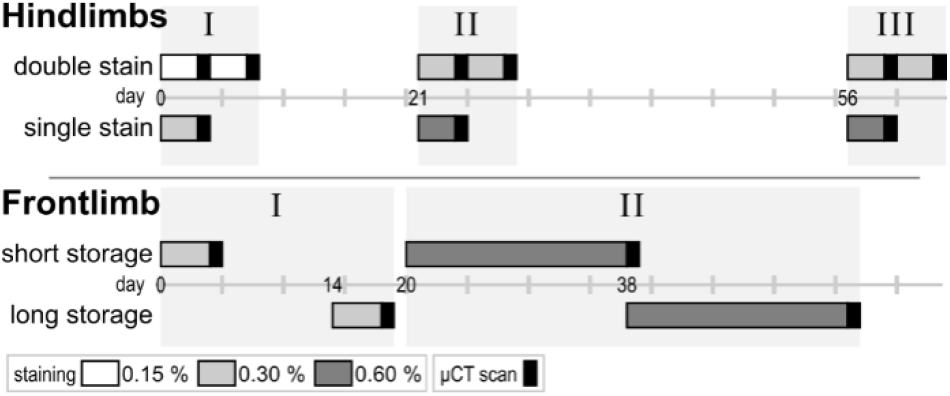
Staining procedure for mouse limbs: hind limbs were stained for 3 days at day 0 with 2 * 0.15% vs. 1 * 0.3% (I) and at day 21 (II) and 56 (III) with 2 * 0.3% vs 1 * 0.6%; front limbs were stained for 4 days with 0.3% at day 0 vs. day 14 (I) and for 18 days with 0.6% at day 20 vs. day 38 (II); n = 8 for all groups

#### Front Limb Staining

In order to determine the effect of the storage time on the final staining result mice front limbs were stained immediately after fixation/rehydration or after various different storage times in 70% ethanol (Figure 1). Therefore the front limbs were separated into a short storage (approximately 1 day) and a long storage (14 days after short storage) group, each containing 4 left and 4 right mouse front limbs. The front limbs were stained using 12.5 ml of a 0.3% Lugol’s solution for 96 h. A 0.3% concentration was selected because it was the concentration used by Metscher [9,10] and is on the lower end of concentrations used in other studies [11]. The long storage group was stored in 70% ethanol and stained 14 days after the short storage group. Both groups were stained a second time with 0.6% Lugol’s solution for 18 days. The long storage group was stained 18 days after the short storage group. A 0.6% concentration was chosen as twice the concentration of the first iteration so that a clearly visible increase in stain in the tissue is achieved.

After the staining duration, the iodine-potassium-iodide solution takes on a yellowish translucent appearance. We assume that this is an indication that the uptake of stain is mostly complete. To minimize any change to the staining by iodine being washed out, the tissues were kept in the residual staining solution until the following staining step. A change in staining is still possible via absorption of the residual Lugol stain and a gradual dispersion of the already absorbed stain inside the tissue, but we believe these changes to be small and similar for all samples. All stainings were performed at room temperature.

#### Nerve, Organ and Tissue Sample Staining

The organ samples were stained the same way as the double stained hind limbs using the same concentrations with 15 ml of Lugol’s solution per cm^3^ of tissue. Tissue volume was measured in the μCT scans of the unstained tissue and organ samples.

The nerve grafts were stained using 0.3% Lugol’s solution for 48 h using approximately 60 ml Lugol’s solution per cm^3^ of sample to achieve a complete staining in a single step. Sample volume was approximated by measuring length and diameter of the graft and the protruding nerve ends and calculating the volume of cylinders with these dimensions.

#### pH Measurement and Density of Residual Staining Solution

To determine the acidity of the staining solution and amount of iodine left in the solution after the staining duration, a pig liver was cut into 10 cm3 sized pieces and stained with 0.3% and 0.6% Lugol’s solution in water or 1xPBS at a ratio of 3 ml solution per cm3 of tissue and stained for 4 days. The samples were stained with a comparatively low amount of solution to ensure a high degree of absorption of the iodine after the staining duration. pH was measured before samples were placed in the solution and daily thereafter with a calibrated Mettler Toledo benchtop pH meter (Mettler Toledo Inc., Columbos, OH, USA).

### Imaging

μCT scans were performed using a SCANCO μCT 50 (SCANCO Medical AG, Brütistellen, Switzerland) specimen μCT scanner.

To avoid movement artifacts, all limbs were placed in 15 ml conical centrifuge tubes in the residual staining solution. Scanning was performed after each staining step with 90 kVp (200 μA, 0.5 mm Al filter, 500 projections, 300 ms integration time), at an isotropic resolution of 34.4 μm. The hind limbs were additionally scanned at an isotropic resolution of 10 μm (90 kVp, 200 μA, 0.5 mm Al filter, 1000 projections, 500 ms integration time) after the final staining step.

The organ and tissue samples (except for nerve) were placed in appropriate centrifuge tubes and scanned at 45 kVp before staining and 70 kVp after each staining step in the residual staining solution. 200 μA, 0.5 mm Al Filter, 500 ms integration time and 500 projections were used for all samples. Due to the different sample diameter, the resolution of the scans was differed between samples. The heart and kidney were scanned at an isotropic resolution of 34.4 μm, the tibialis muscles and the brain at 20 μm and the tendons and sciatic nerves at 10 μm.

The excised silk fibrin conduits with sciatic nerves were scanned first after harvesting and again after Lugol staining with 90 kVp (200 μA, 0.5 mm Al filter, 1000 projections, 500 ms integration time) in the residual staining solution, at an isotropic resolution of 7.4 μm in 1.5 ml Eppendorf centrifugation tubes.

The residual staining solution from the pH measurement of 0.3% and 0.6% in water was filled in 1.5 ml centrifugal tubes along with water, and fresh staining solution with 0.3%, 0.6% and 1% concentrations and scanned with 70 kVp (200 μA 0.5 mm Al Filter, 500 projections, 500 ms integration time) in a single scan.

### Image Processing

μCT scans were exported from the μCT system as reconstructed DICOM stacks. The Stacks were loaded into Amira 6.2 (FEI Visualization Sciences Group, Mérignac, France) and each limb was registered to the first scan of that limb using the register images tool to ease in positioning of measurement regions.

### Measurements

The attenuation was measured in blood, tendon and muscle tissue of the hind limb and in muscle on the front limb. Measurements were performed in the same anatomical region in the flexor digitorum brevis of the leg and the digital flexor muscles of the forelimb. (Figure 2)

**Figure 2:**
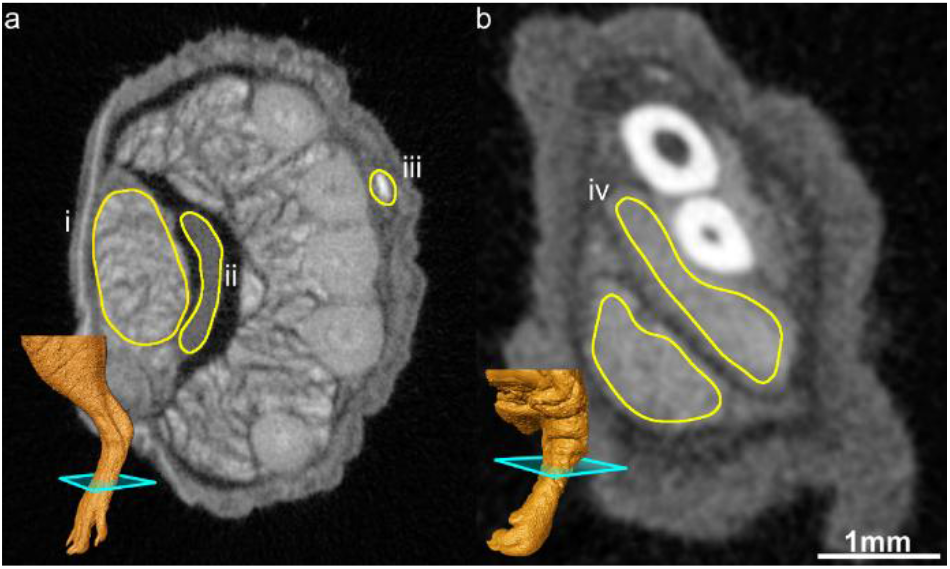
Region of interest (ROI) for attenuation measurements to compare the relative staining of different tissues in close proximity to one another for accurate comparisons; (a) hindlimb ROI in the flexor digitorum brevis (i), tendon (ii) and blood (iii); (b) forelimb ROI in the digital flexors (iv); 3D image in the lower left shows the slice position

The images were imported into Fiji[26,27] and regions of interest (ROI) were drawn by hand using the lasso tool. The average linear attenuation coefficient values were measured in these regions. Due to the small diameter of the blood vessels in relation to the scan resolution, the average intensity is impacted by the partial volume effect[28]. To counteract this, the maximum intensity was also measured in each region, resulting in a measure of blood intensity which is less dependent on the vessel diameter.

The scans of the organ samples at different concentrations were registered to the unstained sample using DataViewer 1.5.6.2 (Bruker microCT, Kontich, Belgium). Using Fiji, the length of the organ samples were measured in four different orientations. The volume shrinkage was calculated from the measured length as the average diameter to the third power.

### Histological Verification of Nerves

After storage in 70% ethanol, dehydration was completed with further increase of a graded series of ethanol up to a concentration of 100%. The nerve samples were then embedded in paraffin *via* the intermedium clearing agent xylol and sectioned at a thickness of 3–4 μm. Samples were then deparaffinized and stained with Martius yellow, Crystal red and Methyl blue (MSB, as described by Lendrum et al. 1962, Studies on the character and staining of fibrin), effectively staining connective tissue blue, fibrin red and erythrocytes yellow, and with Luxol fast blue stain, contrasting myelin.

The histological slides were scanned using an Olympus BX61VS Virtual Slide Scanner (Olympus Corporation, Tokyo, Japan) at a resolution of 0.321 μm/pxl.

μCT scans and histological images were imported into Amira 6.2 (FEI Visualization Sciences Group, Mérignac, France). The position of the histological slide was identified in the μCT scan using a slice object. The slice was manually adjusted to correspond as close as possible to the histological slide.

### Statistical Evaluation

Statistics were calculated using R 3.4.3[29,30]. Error bars in all figures denote the standard deviation. A Student’s t-Test was used for significance testing with a significance level of p < 0.05.

## Results

### Contrast is dependent on total amount of iodine rather than concentration

We think that the small amount of staining solution (15 ml/cm3) combined with the low concentrations (0.15%-0.6%) leads to equilibrium between the tissue and the residual staining solution before the absorption capacity of the tissue is exhausted. The remaining staining solution takes on a yellow translucent appearance in contrast to the opaque brown color beforehand. We hypothesized, that staining at multiple increments at a lower concentration would lead to a similar staining result as staining half as often with double the concentration since the amount iodine accumulated in the tissue would be the same. As shown in Figure 3 it is apparent, with the exception of blood at the highest concentration, that double staining produces a slightly lower attenuation value than single staining. The final attenuation coefficient is very similar however, and does not result in a noticeable change in contrast between different tissues.

**Figure 3:**
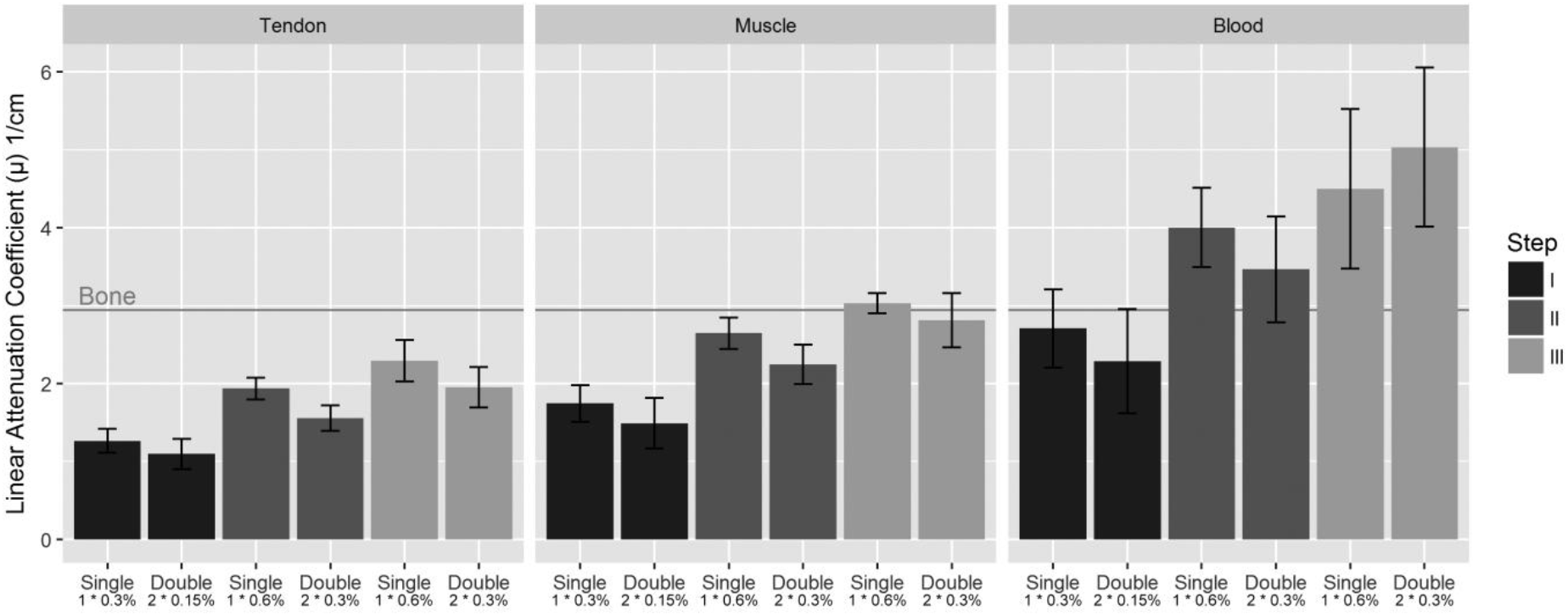
Comparison of linear attenuation coefficient (μ) of different tissues with single vs double staining; Shown are μ values of tendon, muscle and blood tissue with single and double staining with increasing amounts of iodine (I, II and III); n = 8 ±SD; The horizontal line shows the μ value of bone

The contrast can be measured as a ratio of attenuation coefficients between tissue types, which remains mostly unchanged between the concentrations tested and between single and double staining (Figure 4).

**Figure 4:**
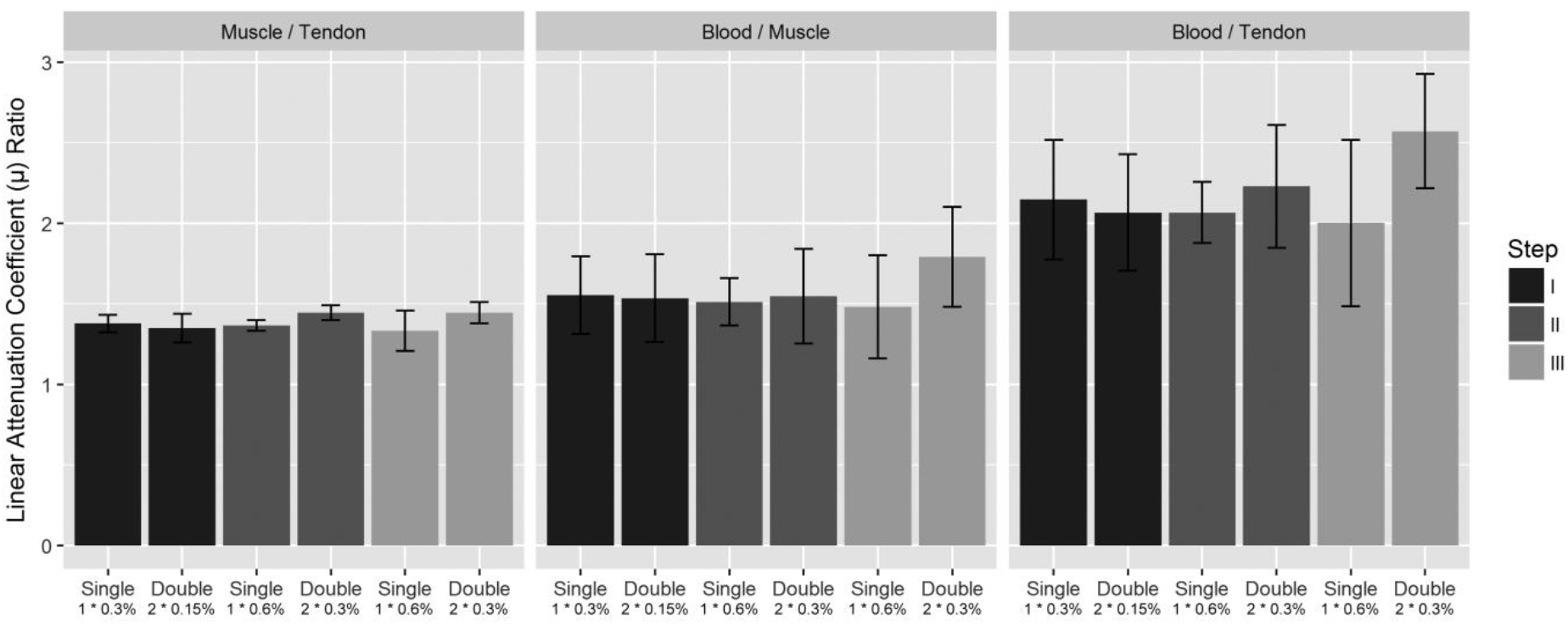
Contrast between tissues at different iodine concentrations with single vs double staining; Shown is the ratio of linear attenuation coefficient (μ) of Muscle / Tendon, Blood / Muscle and Blood / Tendon with single and double staining with increasing amounts of iodine (I, II and III), E.g. Muscle / Tendon single staining at step I have a ratio 1.38, meaning that the muscle exhibits a μ that is 1.38 times higher than in the tendon; n = 8 ±SD

### Storage time negatively influences further stain uptake

The influence of storage time before staining was tested in mice front limbs. No significant difference was observed in muscle tissue when storing the sample in 70% ethanol for an additional 14 days. The front limbs of the short storage group showed an increase in μ value in muscle tissue while there was almost no gain in μ in muscle tissue in the long delay group. A significant difference was observed in the final μ values in muscle tissue between short and long storage before staining (t-Test, p = 0.033) (Figure 5). After the initial staining, thicker parts of the limb were not fully penetrated by the stain. Contrary to the well stained digital flexors in the forelimb, the biceps brachii muscle remained mostly unstained after the first staining step. In the second staining step, the biceps brachii is fully stained but the μ values in this region was also significantly lower after the long storage (t-test, p = 0.012).

**Figure 5:**
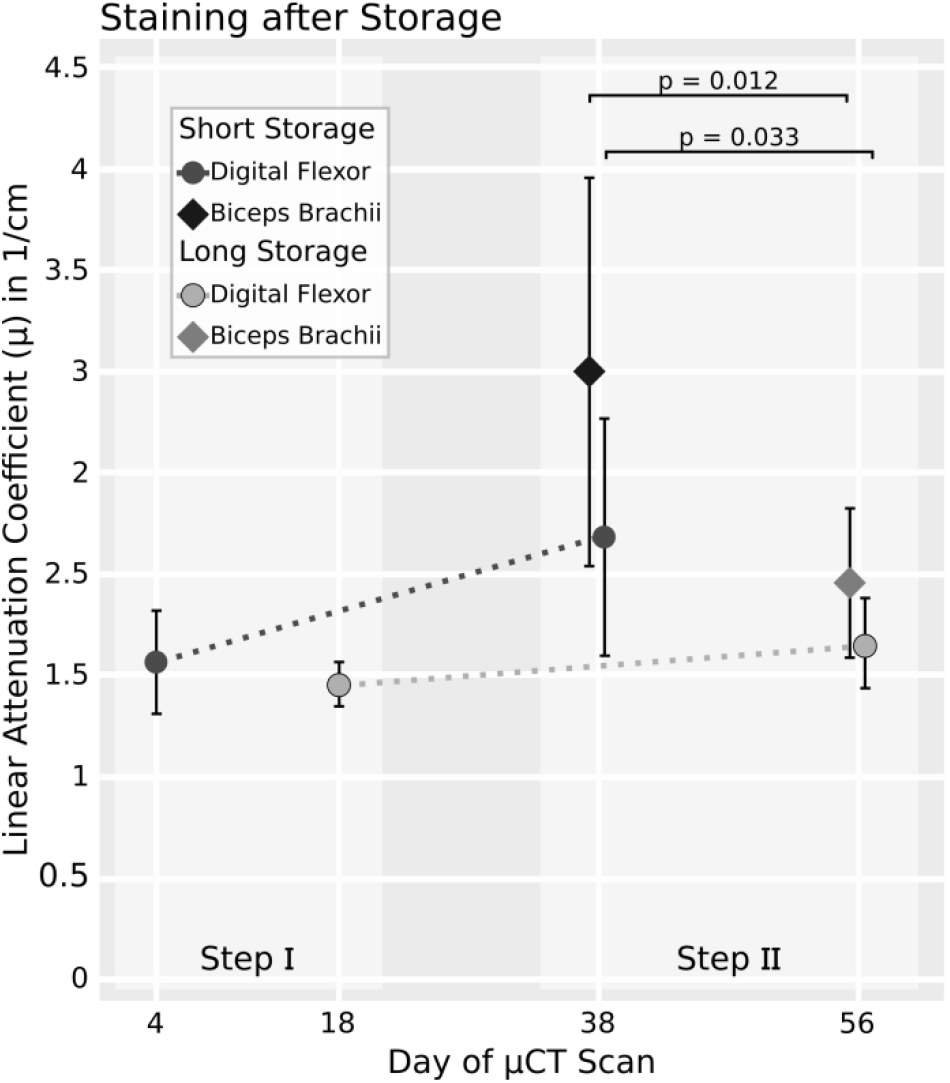
Effect of storage time on muscle tissue staining result; Shown are the linear attenuation coefficients (μ) of the digital flexors and the biceps brachii in the forelimb for the short and long storage time groups at staining steps I (first staining with 0.3%) and II (restained with 0.6%); biceps brachii was not measureable in step I because the stain had not penetrated deep enough; No difference in the staining was observed after step I in the digital felxor. However, significantly lower μ values in both the digital flexors (p = 0.033) and the biceps brachii (p = 0.012) were observed after step II; n = 8 ±SD

### Residual staining solution

Linear attenuation coefficient (μ) measurement of the residual staining solution was performed after 4 days and shows that a considerable amount of stain remains in the solution even after staining is complete. Figure 6 shows a comparison of the μ values of residual staining solutions to the fresh solutions and water.

**Figure 6:**
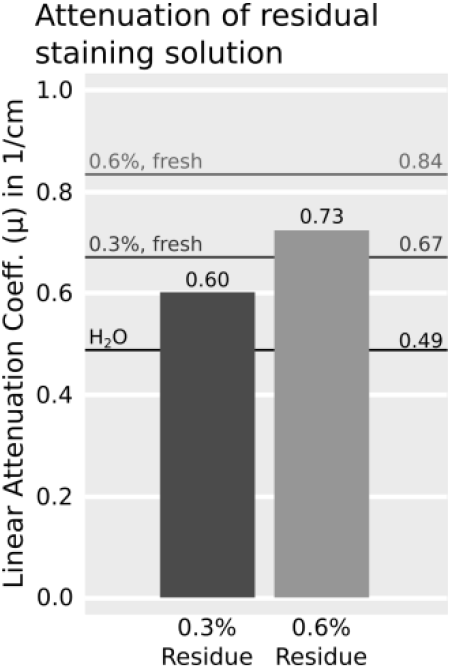
Attenuation of the residual staining solution in comparison to freshly mixed solution and water; after the staining duration the residual solutions become transparent but still retain increased attenuation compared to water

The residual solutions exhibit a translucent appearance and are both much less opaque than the fresh 0.3% solution. Despite the low opacity of the residual solutions, 63% of the increased μ in comparison to water are retained at 0.3% and 68% are retained at 0.6% solution.

### Shrinkage is tissue and concentration dependent

The overall size of the mouse limbs remained the same in our experiments, there was however a widening of the gaps between muscle fascicles and between muscle and other tissues (Figure 7). While individual tissues shrink, the bone provides a rigid support for the soft tissue and thus prevents a size reduction of the whole sample[32].

**Figure 7:**
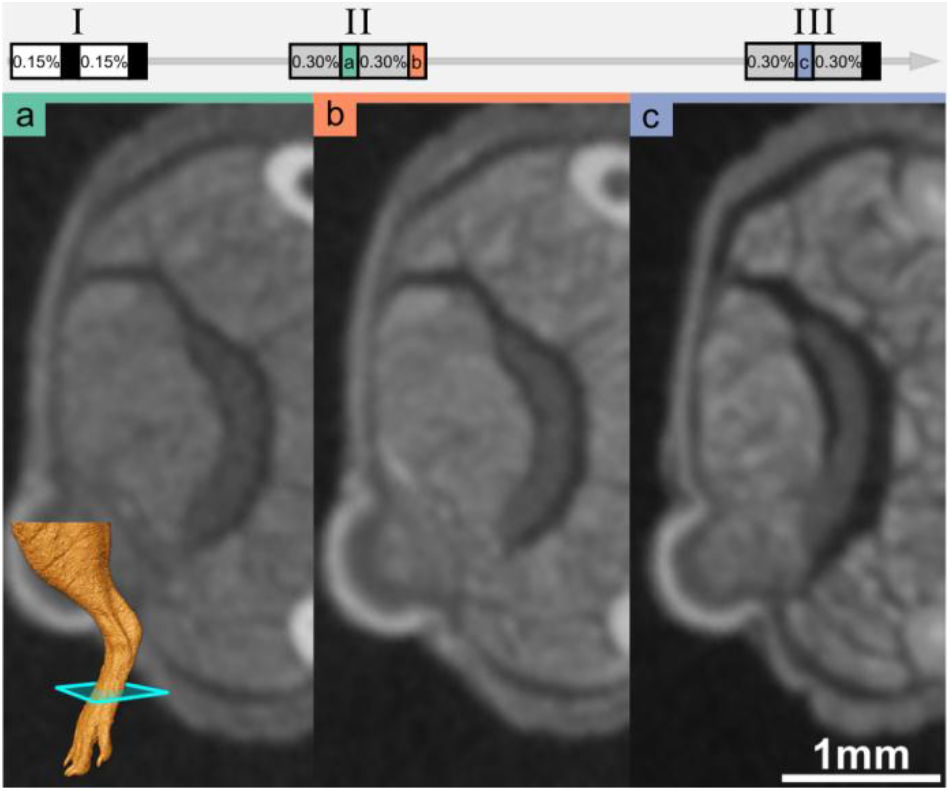
With progressive shrinkage, gaps appear between the muscle fascicles and between different tissues. (a) Staining with 2*0.15% + 1*0.3%: clear gaps between muscles and the tendon, no obvious gaps between fascicles. (b) Staining with 2*0.15% + 2*0.3%: expanded gaps between muscles and tendon. (c) Staining with 2*0.15% + 3*0.3%: gaps between muscles, tendon and skin even wider, gaps between fascicles more pronounced.

All separated organ samples were stained at the same concentrations and the same amount of stain per volume as the hind limbs to determine the shrinkage in different tissue types. With the exception of tendon, which does not appear to shrink due to staining with Lugol’s solution, all tissues shrunk considerably (5-25% change in volume) when being placed in 0.15% Lugol’s solution over a period of 72h (Figure 8). Shrinkage got progressively more severe with consecutive staining iterations (15-43% change in volume after the final staining).

**Figure 8:**
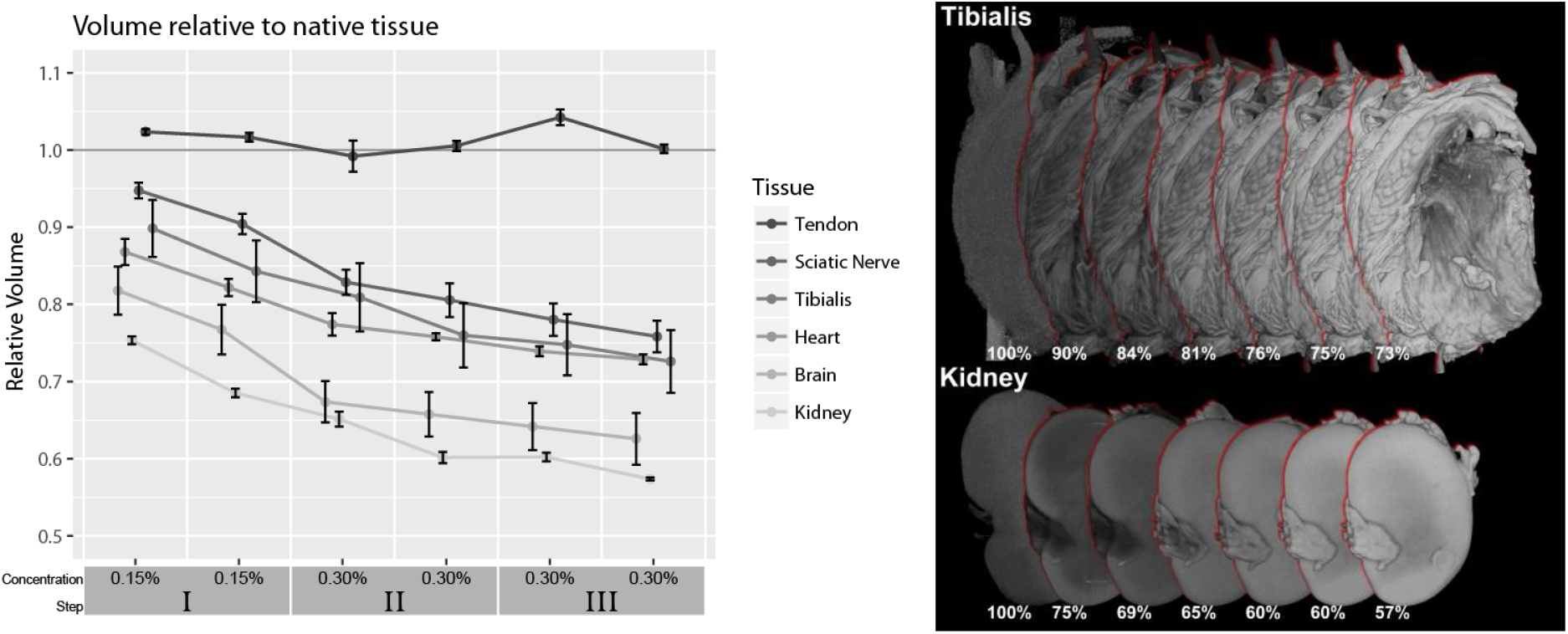
Soft tissue shrinkage at increasing concentrations. Considerable shrinkage already occurs when the samples are initially placed in 0.15% Lugol’s solution. Shrinkage increases with consecutive staining iterations. On the right, volume rendering of μCT images of tibialis and kidney before staining and after each staining step. Percentage values are relative to the volume of the unstained tissue

### Staining homogeneity

With low concentrations or a too short staining period, the iodine does not penetrate all the way into the tissue. This results in a stronger staining of the surface of samples. Bone is less permeable to the stain and encapsulated volumes such as the marrow space are mostly unstained at lower concentrations. We also observed that the foot was less heavily stained than the upper thigh. Where muscle was directly exposed to the staining solution, it was also more heavily stained than muscle protected by skin.

Since most of the iodine in the staining solution is taken up by the tissue at these low concentrations, it is expected that the amount of stain will be a major contributor to the final grayscale value. The fore limb received slightly less stain than the hind limb (~17 ml/cm^3^ vs. ~20 ml/cm3) which may explain the lower attenuation values in the digital flexors of the fore limb compared to the flexor digitorum of the hind limb.

### Decalcification

As described previously, acidic stains such as phosphotungstic acid (PTA) can lead to decalcification of bone tissue and soft tissue shrinkage[9]. We also observed significant decalcification as a result of Lugol staining at high concentrations, indicated by a localized loss of intensity in the μCT scan.

No decalcification was observed in the hind limbs in staining step I which included staining up to a concentration of 0.3%. In staining step II minor decalcification was observed where the bone was directly exposed to the staining solution (Figure 9: b, c) but was not noticeable in bone surrounded by tissue. Starting at staining step III, which repeats the staining of step II (Figure 1), the outer surface of the femur and tibia were decalcified (Figure 9: d, e). Decalcification appears more severe when the bone is adjacent to more heavily stained tissue. Bone which was exposed directly to the staining solution or when it lies directly below the skin was also more severely decalcified.

**Figure 9:**
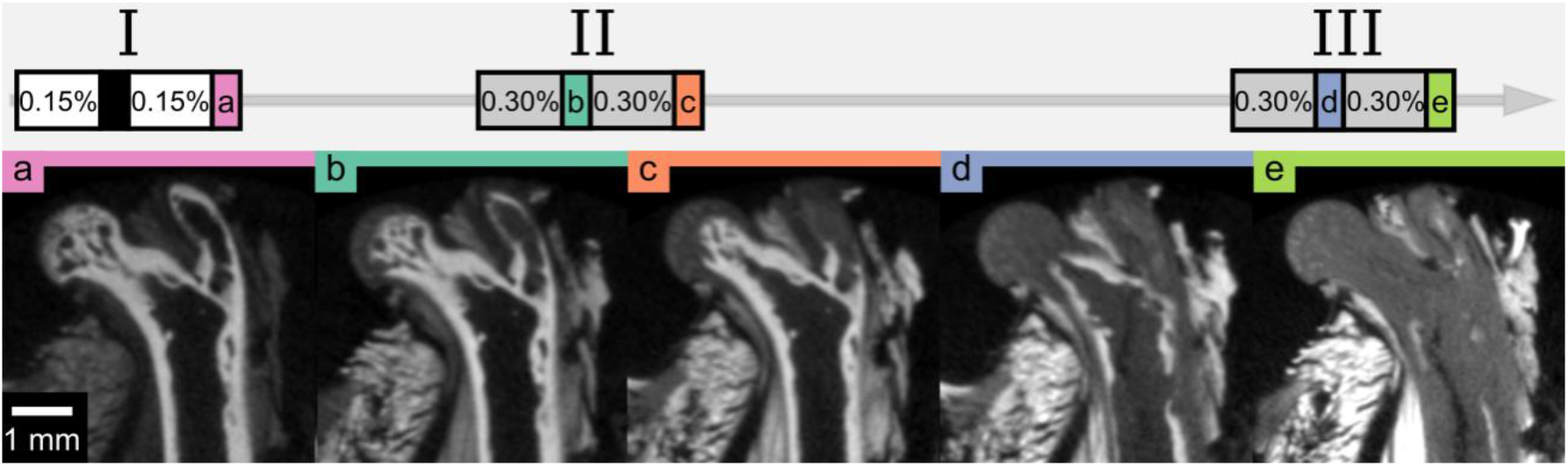
Progressive decalcification during Lugol staining; shown is the femoral head where most severe cases of decalcification were observed. No decalcification was observed in step I (a); in step II exposed parts of the bone exhibit a loss of attenuation indicatng localized decalcification (b, c); in step III larger regions of the bone are affected (d, e)

### Acidity increases during staining

Freshly mixed Lugol solution is weakly acidic depending on the concentration of the solution between pH 4.79 in 0.3% to pH 4.64 in 0.6% in our experiments. The pH begins to decrease further once the staining begins (Figure 10).

**Figure 10:**
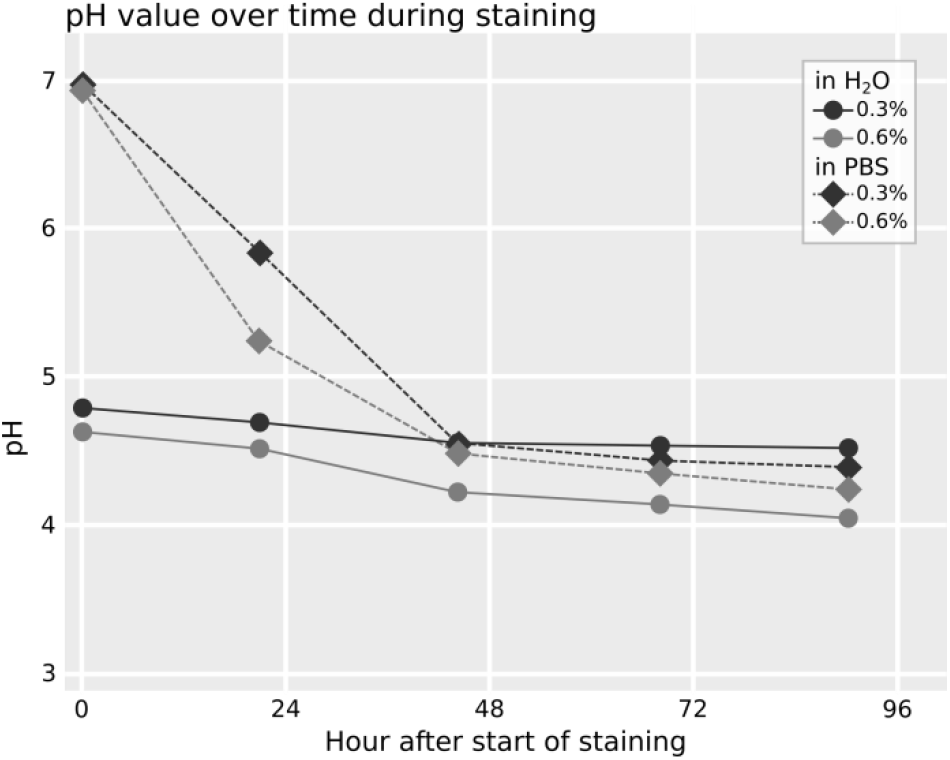
The acidity of the solution is dependent on the concentration of the Lugol solution and increases over time during staining

Using PBS instead of water kept the pH close to 7 before staining but once the staining began the pH value dropped sharply and reached acidity similar to water after 48 hours (pH 4.56 in 0.3% with water vs pH 4.55 in 0.3% in PBS and pH 4.23 in 0.6% with water vs pH 4.49 at 0.6% with PBS).

### Histological verification of nerve regeneration

When staining at a low enough concentration so that the tissues do not get saturated with iodine, the observed gray values depend on the total amount of the stain used. When trying to identify a tissue of interest, the absolute attenuation is thus not a reliable indicator for the type of tissue and different tissues can only be identified by their morphology and in comparison to surrounding tissues. Figure 11 shows a comparison of the histological stainings MSB and Luxol with the corresponding positions in the μCT (Figure 11: c). The silk fibroin conduit is brightly stained by Lugol and has a high intensity in the μCT scan. On MSB stained samples the conduit fibers are visible in red. The position of the histological slice can be found in the μCT based on landmark structures visible in both imaging modalities. In higher magnifications (Figure 11: d), the correspondence between histology and μCT is apparent. Barium sulfate contrast agent visible via histology results in a high attenuation in the μCT scan accordingly (i). A large blood vessel which was not perfused with the perfusion contrast agent is visible because of its low attenuation (ii). When perfusion does not succeed, the unfilled vessels are completely invisible in the μCT without Lugol’s staining. In this regard Lugol’s staining can provide a valuable backup for evaluation of blood vessels. Axonal regeneration is visible in the center of the graft (iii). The myelin of the regenerating nerve, visible in the Luxol stained histology in blue, absorbs more iodine then the surrounding tissue making it also visible in the μCT. On the maximum intensity projection (a), the position of the histology is shown in yellow. The contrast agent perfused blood vessel (i) is also visible here and can be followed in 3D. It can be shown that this blood vessel originates on the outer surface of the graft. It then penetrates the graft proximal to the position of the histological slide and continues in a distal direction where it crosses the position of the histological slide. The myelinated axons (iii) can also be followed allowing the regeneration front to be identified in 3D. On the minimum intensity projection (b) the blood vessel without perfusion contrast agent (ii) can be followed from its proximal origin across the whole graft.

**Figure 11:**
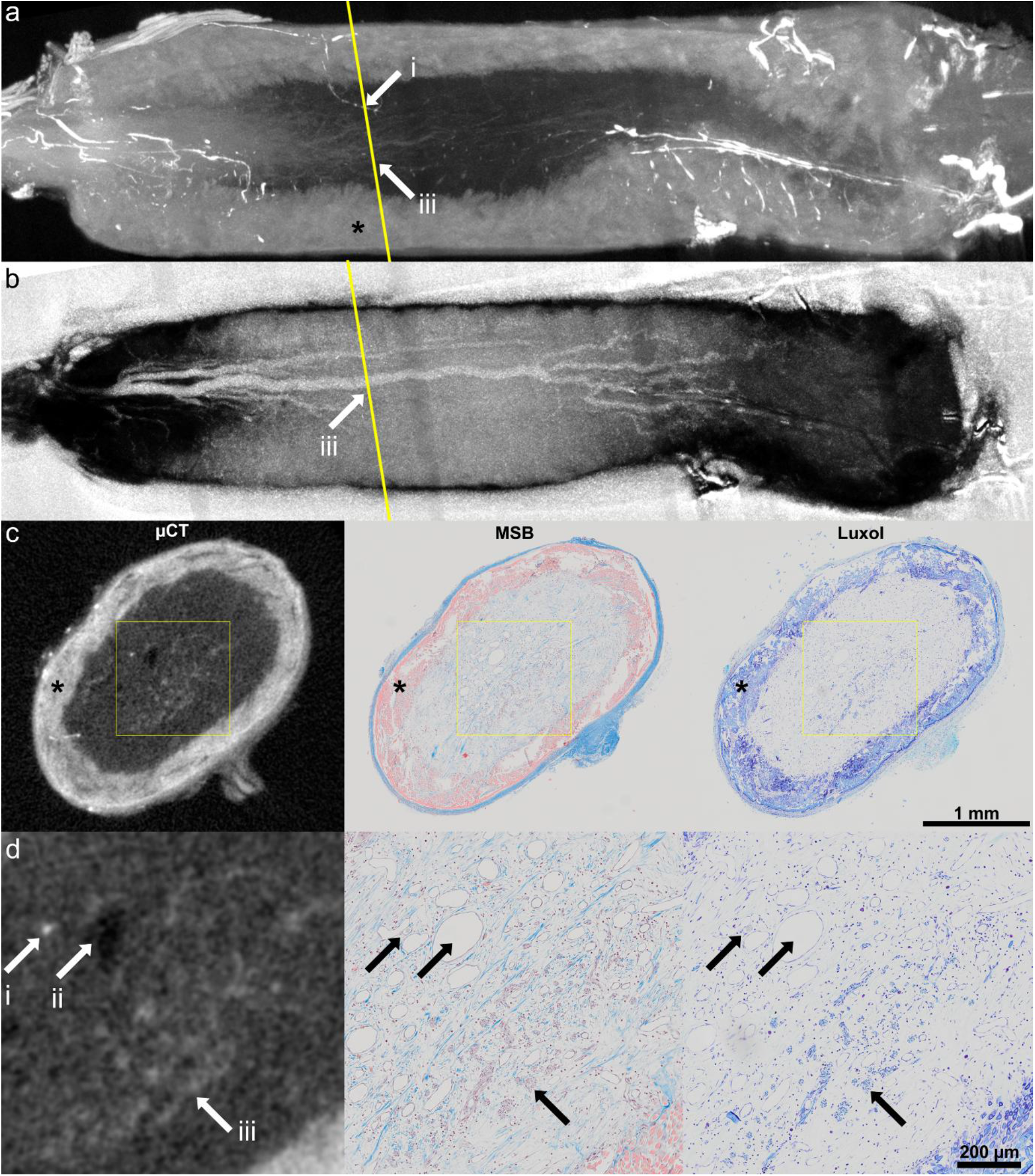
Comparison of μCT with histology work up of a regenerating peripheral nerve in a silk fibroin conduit (asterisk), (a) maximum intensity projection showing the 3D context of perfused blood vessel (i) and regenerated axons (iii), (b) inverted minimum intensity projection visualizes bridging of the defect by blood vessel without perfusion with contrast agent (ii); (c) Position of the histological slide can be found in μCT due to landmark structures, such as blood vessels contrast agent perfused blood vessels (i), blood vessels without contrast agent (ii) and myelinated axons (iii) can be identified in histology and μCT, Arrows i,ii and iii indicate the used landmark structure of the regenerated tissues

## Discussion

In contrast to hard tissue regeneration, which can easily be investigated by CT based imaging, the field of soft tissue regeneration lacks a high resolution 3D imaging method. For the analysis of soft tissue regeneration by μCT the main technical requirement is a reliable staining of the tissue of interest while minimizing deleterious influences on the morphology of the tissue. We could show that a good contrast between different tissues can be achieved at different iodine concentrations and while higher concentrations stain tissue quicker, the contrast between tissues remains mostly the same with the amounts of stain used. Storage in ethanol for several days before staining does appear to reduce the uptake of stain into the tissue (Figure 5). Differences in attenuation were only observed after the second staining step. This suggests that the tissue still takes up stain at a comparable rate after storage but with a reduced uptake capacity. However, long-time storage in ethanol is not recommended as the occurring delipidation limits the overall contrast differences between tissues with different lipid densities[11]. The hind limbs were stored in Lugol solution a comparable time to the forelimbs between staining steps with no obvious negative effects without any apparent loss in contrast between tissues (Figure 4). This suggests that consistency is a more important factor in producing comparable stains than the absolute storage time. Nonetheless, when comparing different samples, we would advise to keep any delays as short as possible and the same for all samples to minimize decalcification and any other possible impacts on the staining result. Since the limbs were scanned in the residual staining solution, the attenuation of the residual solution may have an impact on the measured attenuation. The double staing group of the hind limbs was always stained at half the concentration of the single staining group. This difference in concentration would result in a slight difference in attenuation of the residual staining solution which may be partially responsible for the slightly lower attenuation of the double staining group (Figure 3). However, the difference in attenuation between the residual solution is small compared to the attenuation of the tissue (Figure 6) and differences are threfore expected to be minor.

When staining with a low enough concentration of Lugol’s solution, we believe that iodine is absorbed by the tissue until equilibrium is reached between the amount of iodine in the tissue and the staining solution. The final attenuation value in the tissue is determined by the amount of iodine absorbed by the tissue. Since, at the concentrations used, the absorption capacity of the tissue is not fully exhausted, not only the concentration but also the absolute amount of stain has to be considered. We selected fore- and hindlimbs because they not only have a very consistent size, they also have a range of different tissue types at almost identical relative volume. When planning staining of a tissue sample, the total volume and the relative volume as well as the different affinity towards the stain influence the staining result. Lugol’s solution has been reported to preferentially stain glycogen[33,34] which has an exceptionally high binding capacity to iodine[35]. Moreover, iodine binds to lipids, such as found in myelinated nervous tissue[36].

Considering that the exact makeup of the sample is often unknown, developing a suitable staining protocol can therefore be challenging.

As observed by Buytaert et. al.[31], tissue shrinkage does occur in soft tissue with iodine staining at higher concentrations. We have observed that tissue shrinkage is increased at higher amounts of stain (Figure 8). Since the volume of the organ samples decreases between the two stainings in each staining step, which are stained at the same concentrations, the shrinkage appears to be dependent on the total amount of iodine in the tissue, rather than the concentration of the staining solution. Shrinkage appears to be unavoidable, considering that considerable shrinkage occurs already in the first staining step which contains too little iodine to fully stain any of the organ samples with the exception of the sciatic nerves. Similar results were observed by Vickerton et. al. at higher iodine concentrations (2-20%), also recommending using the lowest amount of stain necessary to achieve good contrast. Unfortunately, they do not provide the total amount of staining solution used or whether it was adjusted to the sample volume[37]. Wong et. al.[38] developed a method of reducing shrinkage by stabilizing the tissue in a hydrogel, observing 20% difference in volume when comparing stabilized and un-stabilized mouse embryo samples. To completely avoid shrinkage, other stains such as Hafnium-based Wells-Dawson polyoxometalate (Hf-POM) can be considered which does not appear to result in any shrinkage[39] and was shown to be a promising contrast agent for measurements in soft tissue in μCT[40]. However these stains are rather expensive in their production and stain different structures.

Decalcification in bone regions not directly exposed to the staining solution only appeared at staining Stage III (corresponding to a cumulative amount of 1*0.3%+2*0.6% or 2*0.15%+4*0.3% respectively), at which point the bone had been exposed to the stain for 60 days, possibly due to the weakly acidic nature of Lugol’s solution (Figure 9). No decalcification was observed in the foot, where the staining appears generally less intense than in the calf or thigh. In the front limbs, decalcification also occurred despite the lower concentrated Lugol’s solution. This suggests that decalcification increases in severity with longer staining times. The decalcifying effects of Lugol staining are rarely discussed in the literature. Gignac et. al.[36] adapts Lugol staining to larger samples, requiring longer staining (up to 4 weeks) in higher concentration (11.25%) than commonly used, yet does not mention decalcification. Sayin et. al.[41] analyzed the decalcifying effects of 2% Lugol’s solution as an endodontic irrigant and found a significant increase of Ca^2+^ ions when exposing the dentin of a human root canal to the Lugol’s solution compared to distilled water, thus confirming the decalcifying properties. The decalcifying effect of Lugol’s solution appears to be relatively minor even compared to other endodontic irrigants, exhibiting a Ca^2+^ release about four times lower than 2.5% NaOCl[41], the decalcifying properties of which have previously been shown in a similar setup to be almost an order of magnitude lower than 15% ethylenediaminetetraacetic acid (EDTA)[42]. Lugol staining is usually employed to make non-calcified tissues visible in the μCT which, combined with the relatively minor decalcifying effect, may explain why the decalcification is so rarely observed and discussed. However, if calcified structures are of interest, the decalcifying effect of Lugol’s solution should be considered.

Tissue penetration is faster at higher concentrations. A short staining time is desirable both for a quicker turnover but also to minimize negative effects such as shrinkage and decalcification of the bone tissue. On the other hand, higher concentrations which shorten the staining time also lead to an increase in tissue shrinkage. For anatomical applications several studies exist where staining parameters are suggested, many of which are listed in Gignac et al.[11]. However for tissue regeneration purposes where the makeup of the tissue differs strongly from the physiological case and can be further complicated by the presence of non-site-specific materials such as tissue grafts or synthetic implants, such references can provide only limited help. To find the ideal parameters, individual testing appears unavoidable.

Overstaining the tissue increases the negative effects of the staining and should be avoided. We have however shown that applying lower concentrations in multiple steps can result in a very similar outcome to staining in a single high-concentration step. So even if preliminary experiments to optimize the staining are not providing the desired result immediately, a good result can still be achieved with a stepwise staining while avoiding overstaining. Li et. al.[43] suggests a stepwise staining approach with increasing concentrations to preserve a similar concentration difference between the staining solution and the tissue. They observed a more homogenous distribution of stain within the tissue with less shrinkage compared to staining in a single step but with an increased staining duration using their stepwise staining approach.[43]

In anatomical studies, different tissues can often be identified by their location and morphology[9]. When the nature of a tissue is uncertain, correlative histological verification can be used to directly compare the same site with a histological image where the tissue can be clearly identified[44]. In the field of tissue regeneration, where new tissue grows, a proper identification of the tissues of interest is crucial. In case of the sciatic nerve grafts, we could show that regenerating nerve fibers can be identified in the μCT scans. This was confirmed by registering the scans with the histological images allowing for a direct comparison. In the study for which the sciatic nerve defects were scanned, the penetration of blood vessels through the silk graft was of interest. With the combination of barium sulfate perfusion and Lugol staining we could locate blood vessels penetrating the graft. The staining also proved useful for verifying whether or not the defect was fully bridged on samples where the longitudinal cuts missed the center of the regenerating nerve. The correlative imaging approach can aid in an improved understanding of the regeneration processes by providing a valuable 3D context to the histological analysis. The μCT scans can also be used to identify potential regions of interest so that histological sectioning is done at the most interesting positions in the sample.

The Lugol staining had no apparent detrimental impact on the subsequent histological work-up and (immuno-) histological stainings (MSB and Luxol staining, Figure 11, as well as HE, Neurofilament, Glial fibrillary acidic protein (GFAP), data not shown). It should be noted that in combination with sodium thiosulfate, which is routinely used to remove Lugol’s solution from the tissue in clinical settings, a negative effect on CD30 immunostaining after Lugol staining has been reported[45]. Although our protocol does not include a rinsing step and no negative effects have been observed in our lab, we advise to perform preliminary staining tests to verify that immunoreactivity is intact.

We believe that the usage of contrast enhanced soft tissue μCT and its 3D information is able to provide novel insights in a variety of fields. Besides regeneration of peripheral nervous tissue shown in this study, events after central nervous injuries such as spinal cord injury or brain injury could be analyzed.

## Conclusion

Here we present evidence that Lugol staining can provide valuable additional information for understanding tissue regeneration processes. We discuss potential pitfalls of contrast enhanced soft tissue μCT imaging and provide solutions for the usage of Lugol’s solution. We consider individual adjustments to the staining process to be necessary for staining different tissues. Incremental staining can help ease the optimization process, reduce the risk of overstaining and help minimize shrinkage associated with higher concentrations. In this work we are the first to visualize regenerating nervous tissue using contrast enhanced μCT. One of the major advantages of this imaging method is the possibility to identify and preselect sites of interest for subsequent histological sectioning using μCT analysis. Histologic techniques can be used to verify structures visible in the μCT to reduce the risk of misidentifying tissues while the μCT can provide valuable 3D context to histological observations. Correlative imaging of Lugol stained μCT and conventional histology can mutually improve the interpretation in both imaging techniques.

## Data availability

Original μCT data of the hind limbs, fore limbs and organ samples can be made available upon request to the corresponding author.

## Conflicts of Interest

None.

## Funding Statement

The authors received no specific funding for this work.

## Author Contributions

PH, NS, PS, MK, DH, HR participated in the study design. PH, NS, MK, DH Conducted the study. NS, MK, DH performed the surgeries. AT developed and provided the nerve grafts. PH, NS performed μCT scanning. SN performed histology. PH performed the data analysis and wrote the manuscript. PH, AT, DH, PS participated in the data interpretation. PS, HR, AT, DH aided in the review process and provided scientific input.

